# *KmerAperture*: Retaining *k*-mer synteny for alignment-free extraction of core and accessory differences between bacterial genomes

**DOI:** 10.1101/2022.10.12.511870

**Authors:** Matthew P. Moore, Mirjam Laager, Paolo Ribeca, Xavier Didelot

## Abstract

By decomposing genome sequences into *k*-mers, it is possible to estimate genome differences without alignment. Techniques such as *k-*mer minimisers (MinHash), have been developed and are often accurate approximations of distances based on full *k*-mer sets. These and other alignment-free methods avoid the large temporal and computational expense of alignment or mapping. However, these *k*-mer set comparisons are not entirely accurate within-species and can be completely inaccurate within-lineage. This is due, in part, to their inability to distinguish core polymorphism from accessory differences. Here we present a new approach, *KmerAperture*, which uses information on the *k*-mer relative genomic positions to determine the type of polymorphism causing differences in *k*-mer presence and absence between pairs of genomes. Single SNPs are expected to result in contiguous series of relative unique *k*-mers of length *L*= *k*. On the other hand, series of length *L* > *k* may be caused by accessory differences of length *L*-*k*+1; when the start and end of the sequence are contiguous with homologous sequence. Alternatively, they may be caused by multiple SNPs within *k* bp from each other and *KmerAperture* can determine whether that is the case. To demonstrate use cases *KmerAperture* was benchmarked using datasets including a very low diversity simulated population with accessory content independent from the number of SNPs, a simulated population were SNPs are spatially dense, a moderately diverse real cluster of genomes (*Escherichia coli* ST1193) with a large accessory genome and a low diversity real genome cluster (*Salmonella* Typhimurium ST34). We show that *KmerAperture* can accurately distinguish both core and accessory sequence diversity without alignment, outperforming other *k*-mer based tools.

## INTRODUCTION

The increasing availability of whole genome sequencing provides us with an opportunity to transform our analytical approach to bacterial population genetics and infectious disease epidemiology^1,2^. This potential for genomic epidemiology is being investigated by public health organisations, such as the United Kingdom Health Security Agency (UKHSA), where routine surveillance involves whole genome sequencing, typing and detailed comparative and phylogenetic analysis^3^. However, once collected and sequenced, the comparison of thousands of bacterial genomes poses a number of important difficulties and limitations, particularly the computational time and expense required to compute alignments. For example, the exhaustive comparison of all pairs within a set of 5,000 genomes would require >12×10^6^ pairwise alignments. This is often shortcut to 5,000 alignments by aligning each genome to a single reference genome and then comparing the genomes at each reference position^1,4^. This is much more efficient than comparing all pairs of genomes, but it still involves the computation of a large number of alignments. Furthermore, it can introduce a bias depending on which reference genome is used, and also loses any information about accessory regions not found in the reference.

A number of efficient alignment-free approaches have been developed to circumvent these issues. The most efficient of these involve decomposition of genomes into *k*-mers (*k*-mers/kmers/*n*-grams; unique sequences of length *k*), subsampling into ‘sketches’ and performing set analysis^5–8^. Methods such as the min-wise independent permutations local sensitivity hashing scheme (MinHash) algorithm, accurately approximate the full-set differences across genomes and are extremely fast^5^. However, the accuracy in estimating relatedness decreases the more closely related the genomes are for full *k*-mer set and MinHash distances. Alignment-free distance estimates tend to correlate primarily with quantities of differentially present sequences rather than SNP distances. For closely related genomes, up to the point of re-sequenced isolates, this won’t by definition correlate with polymorphism^9^.

Highly similar genomes are those that are most relevant for outbreak detection and further analysis for public health^10–12^. For instance, MinHash-based distance measures will not accurately discern clusters at a lower diversity level than an MLST-defined lineage. Ultimately, this is due to the inability to know whether mismatching *k*-mers result from polymorphism or relative presence/absence of the sequence. PopPUNK is one approach that uses MinHash for within-species differences by modelling the change in mismatches across different values of *k* in an attempt to differentiate diversity that is core to the population and diversity that is accessory^13^.

Two approaches have demonstrated an ability to estimate some within-lineage SNPs^14^: kSNP3.0^15^ and Split K-mer Analysis (SKA)^16^, with similar benchmark performance^14,16^. These approaches successfully identify what, in set analysis-based methods, would be represented by mismatched *k*-mers resulting from SNPs indistinguishable from those resulting from accessory sequence.

Here, we present *KmerAperture*, a novel alignment-free algorithm with the ability to determine bacterial genetic differences at scale. Mismatching *k*-mers are linked to their original synteny, taking advantage of initial efficient *k*-mer set analysis to generate the relative *k*-mer complements. Contiguous *k*-mer series may then be investigated as to whether they’re generated by presence/absence of sequences, SNPs or repeats. Crucially, *KmerAperture* is also able to gather SNPs within *k* of one another on the genome, usually a confounding feature of *k*-mer comparisons.

## RESULTS

### An illustrative example

Figure 1 provides a toy example of the *KmerAperture* algorithm compared with full set Jaccard similarity estimates. The reference sequence and query are separated by a single SNP and a further 5bp sequence present only in the query. The sequences are decomposed into *k*-mers where *k* =3. When there is an isolated unshared sequence of length *L* we can anticipate this to produce *L-k+*1 unique *k*-mers covering the unshared sequence. In the case, as in Figure 1, that the unshared sequence is also contiguous with shared sequence it will generate a further *k-*1 unique *k*-mers for each contiguous side. Alternatively, we anticipate a SNP to generate *k* unique *k*-mers in each set except when the SNP is within *k* of another SNP or sequence start or end (such as the end of a contig).

**Figure 1.**
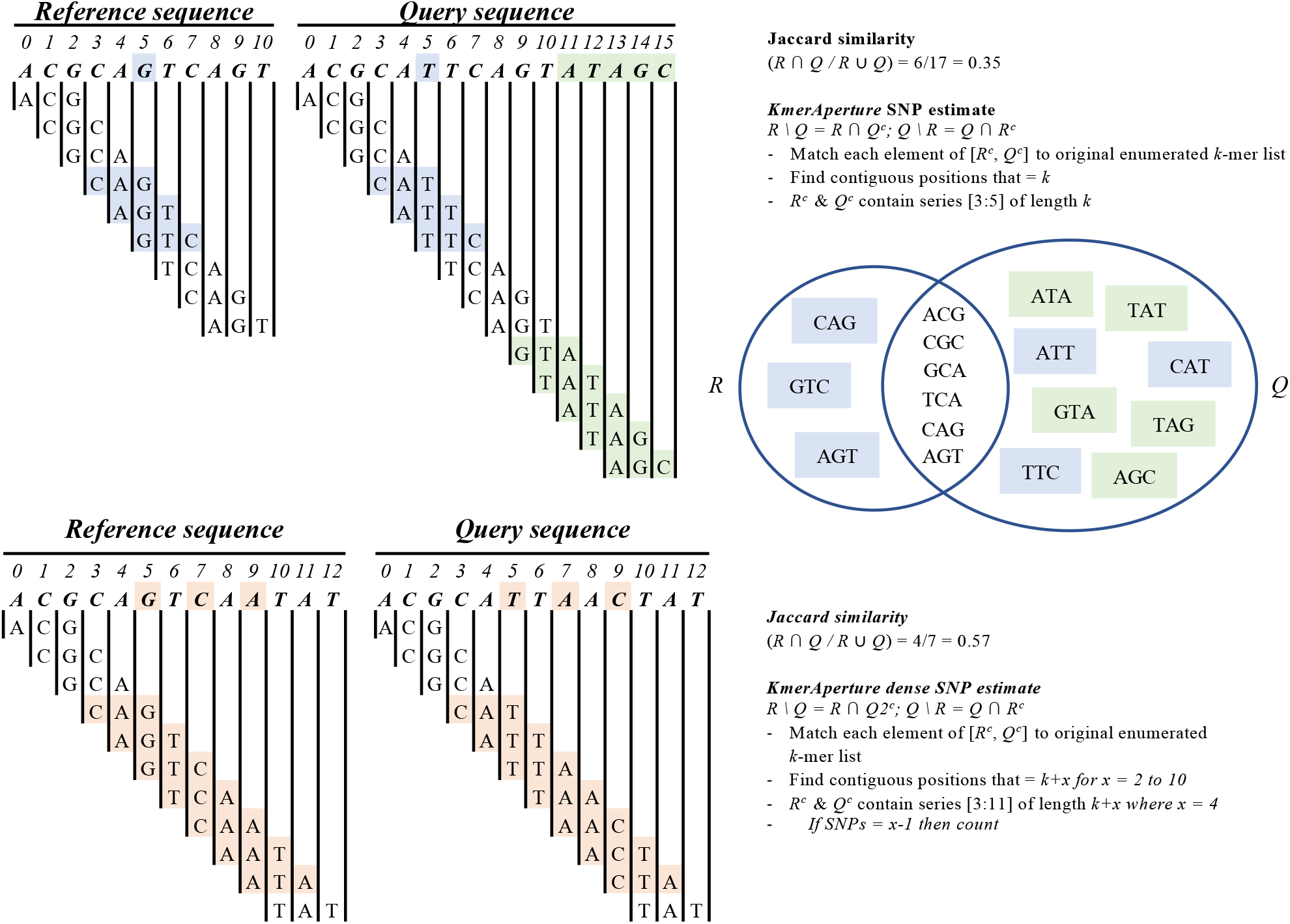
(Upper) Blue shading highlights unique *k*-mers generated as the result of a SNP and green shading indicates unique *k*-mers as the result of an accessory sequence. In the Jaccard similarity these types of mismatches are not differentiated. *KmerAperture* starts by considering the genomic position of the elements unique to *R and Q* respectively. As the blue shaded *k*-mer series equals *k* in length, this would be assessed as to whether the series is comprised of overlapping *k*-mers. Then, the middle *k*-mer with the middle base removed matched between reference and query *k*-mer series that equal *k* in length. In contrast, the green shading demonstrates how *k*-mer series of >*k* may be generated by accessory sequences. (Lower) This example shows with orange shading 3 SNPs where the positions are within *k. KmerAperture* would correctly identify these as all possible pairs of all series of the same length between length *k* and 100*k* would be reconstructed into their original sequence and mismatches counted. This example, which is illustrative, would not however pass the sequence diversity filter of <25%

In Figure 1, *KmerAperture* would correctly discern mismatched *k*-mers resulting from SNPs vs accessory sequence. Both set complements would result in contiguous *k*-mers of serial length *k*. As a result, the mean estimated SNPs would be calculated and it would be determined that in the comparable sequence regions there is a single SNP. *KmerAperture* would additionally discern, from the five contiguous *k*-mers unique to the query sequence, that there was an accessory difference of 5bp (since these *k*-mers are contiguous with shared sequence on one side, they are expected to create (*L*-*k*+1)+(*k*-1)=*L k*-mers). For SNPs within *k* of another on the genome (Figure 1), they will produce series of length *L*>*k*+2 up to a maximum of *S*(*k*-1)+1 where *S* is the number of SNPs within *k* of one another.

### Rationale for benchmarks

To demonstrate its usefulness in a number of relevant use cases, we benchmarked *KmerAperture* in four scenarios: a very low diversity simulated population with accessory content independent from the number of SNPs; a simulated population were SNPs are spatially dense; a moderately diverse real cluster of genomes (*Escherichia coli* ST1193) with a large accessory genome; and a low diversity real genome cluster (*Salmonella* Typhimurium ST34).

### Application to simulated genomes with randomly distributed SNPs

The first simulated genome set includes 500 genomes with varying degrees of accessory difference to the reference and a range of 1-100 SNPs (median 11 SNPs). *KmerAperture* estimated a median (range) of 11 (0-100) SNPs with *k*=19, 25 and 31. SKA produced the same median (range) of 11 (0-100) across all sizes of *k*.

The ability of *KmerAperture* and SKA to identify SNP counts less than or equal to various thresholds was assessed (Table 1). *KmerAperture* produced high positive predictive values and specificity across SNP thresholds of ≤25,20,15,10 and 5. Across all choices of *k* and all thresholds with *KmerAperture* there were no false negatives (no overprediction of SNP values above a threshold), meaning sensitivity was 100% in all cases. There was a small number of false positives with between 2 and 5 for *k*=19 and 1-3 for *k*=31. With SKA there was a similarly small number of false positives with a maximum of 4 SNPs at *k*=19 and a threshold of ≤15 SNPs. With SKA the specificity and PPV was equivalent or greater in all cases. However, there was a single false negative with *k*=19 and 25 for the ≤5 SNP threshold.

**Table 1.** Performance of *KmerAperture* and SKA for identifying SNP distances below a threshold. True positives (*TP*) are those correctly found to be below the SNP thresholds (25, 20, 15, 10, 5), false positives (*FP*) those incorrectly found to be below the threshold, and true negatives (*TN*) those correctly found to be above the threshold. Performance was measured on specificity *TN*/(*TN*+*FP*) and positive predictive value (PPV) *TP*/(*TP*+*FP*)

| Algorithm | *k*-mer size | Accuracy | SNP thresholds |  |  |  |  |
| --- | --- | --- | --- | --- | --- | --- | --- |
| | | | $\leq 25$<br>(*n*=340) | $\leq 20$<br>(*n*=316) | $\leq 15$<br>(*n*=289) | $\leq 10$<br>(*n*=242) | $\leq 5$<br>(*n*=178) |
| *KmerAperture* | *k*=19 | *Specificity* | 0.939 | 0.947 | 0.945 | 0.969 | 0.988 |
|  |  | *PPV* | 0.986 | 0.984 | 0.98 | 0.984 | 0.989 |
|  | *k*=25 | *Specificity* | 0.988 | 0.989 | 0.963 | 0.992 | 0.988 |
|  |  | *PPV* | 0.997 | 0.997 | 0.986 | 0.996 | 0.989 |
|  | *k*=31 | *Specificity* | 0.988 | 0.989 | 0.972 | 0.992 | 0.988 |
|  |  | *PPV* | 0.997 | 0.997 | 0.99 | 0.996 | 0.989 |
| SKA | *k*=19 | *Specificity* | 0.988 | 0.978 | 0.963 | 0.992 | 0.988 |
|  |  | *PPV* | 0.997 | 0.994 | 0.986 | 0.996 | 0.989 |
|  | *k*=25 | *Specificity* | 1.0 | 0.989 | 0.972 | 0.992 | 0.994 |
|  |  | *PPV* | 1.0 | 0.997 | 0.99 | 0.996 | 0.994 |
|  | *k*=31 | *Specificity* | 1.0 | 0.989 | 0.972 | 0.992 | 0.994 |
|  |  | *PPV* | 1.0 | 0.997 | 0.99 | 0.994 | 0.994 |

For *k*=19, 25 and 31, *KmerAperture* estimated the accessory difference for each genome compared with the reference. Relative presence was estimated for each genome in each pair. In all cases, it was correctly estimated that there was no relative additional sequence in the reference genome (0bp estimated). The relative additional query sequence ranged from a total size of 4,139bp to 1,926,407bp with a median of 736,708. This included 169 genomes with >1Mbp additional sequence. *KmerAperture* also attempts to identify these regions and extract them. The median absolute error as a percent of bp different to the real accessory size was 0.0017%, with a maximum error of 27.9% (of 827,256bp accessory).

### Application to simulated genomes with variably densely distributed SNPs

In the following, we define *k*-*clustered SNPs* as chains of SNPs such that each SNP belonging to the chain, within a genome, has at least one neighbouring SNP at ≤*k* nucleotides of distance.

Three genome sets were simulated with 50, 100 and 150 SNPs respectively (n=750 genomes total). Each set contained 250 genomes evenly divided (n=50 genomes) into those with 20%, 40%, 60%, 80% and 100% of their SNPs being clustered within *k*=25 of one another. It was randomly determined how many of the SNPs would form these SNP chains, for instance 50 SNPs being 100% *k*-clustered could involve three *k*-clusters of size 10, 10 and 30 SNPs within *k* of on another. Detecting *k*-clusters is a major weakness of *k*-mer based methods and we expect a degree of non-retrieval in these test conditions. Performance was better for *KmerAperture* than for SKA in the retrieval of SNPs and *k*-clustered SNPs, across all densities (20%-100%) and overall SNP number (50, 100 and 150 SNPs).

The median reduction in retrieval performance with increasing *k*-clustering was smaller for *KmerAperture*. There was a relative decrease of SNPs retrieved from 20% to 100% fraction of 2.78%, 10.95% and 1.8% for 50, 100 and 150 SNPs respectively. For comparison, the relative reduction in performance with SKA between 20% and 100% of *k*-clustered SNPs was 48.48%, 55.97% and 53% for 50, 100 and 150 SNPs respectively. The range in performance increased for *KmerAperture* with increased *k*-clustering with minimum retrieval as low as 8/50 SNPs and 20/100 SNPs retrieved at 100% *k*-clustering and 48/150 SNPs retrieved at 80% *k*-clustering. There was a greater range in performance with *KmerAperture*, however and SKA had a greater minimum retrieval in two categories (50 SNPs, 100% *k*-clustering and 100 SNPs, 20% *k*-clustering). In the remaining 13/15 the greater minimum retrieval was with *KmerAperture*. In all categories the median and max retrieval was with *KmerAperture* (Table 2, Figure 2).

**Table 2.**
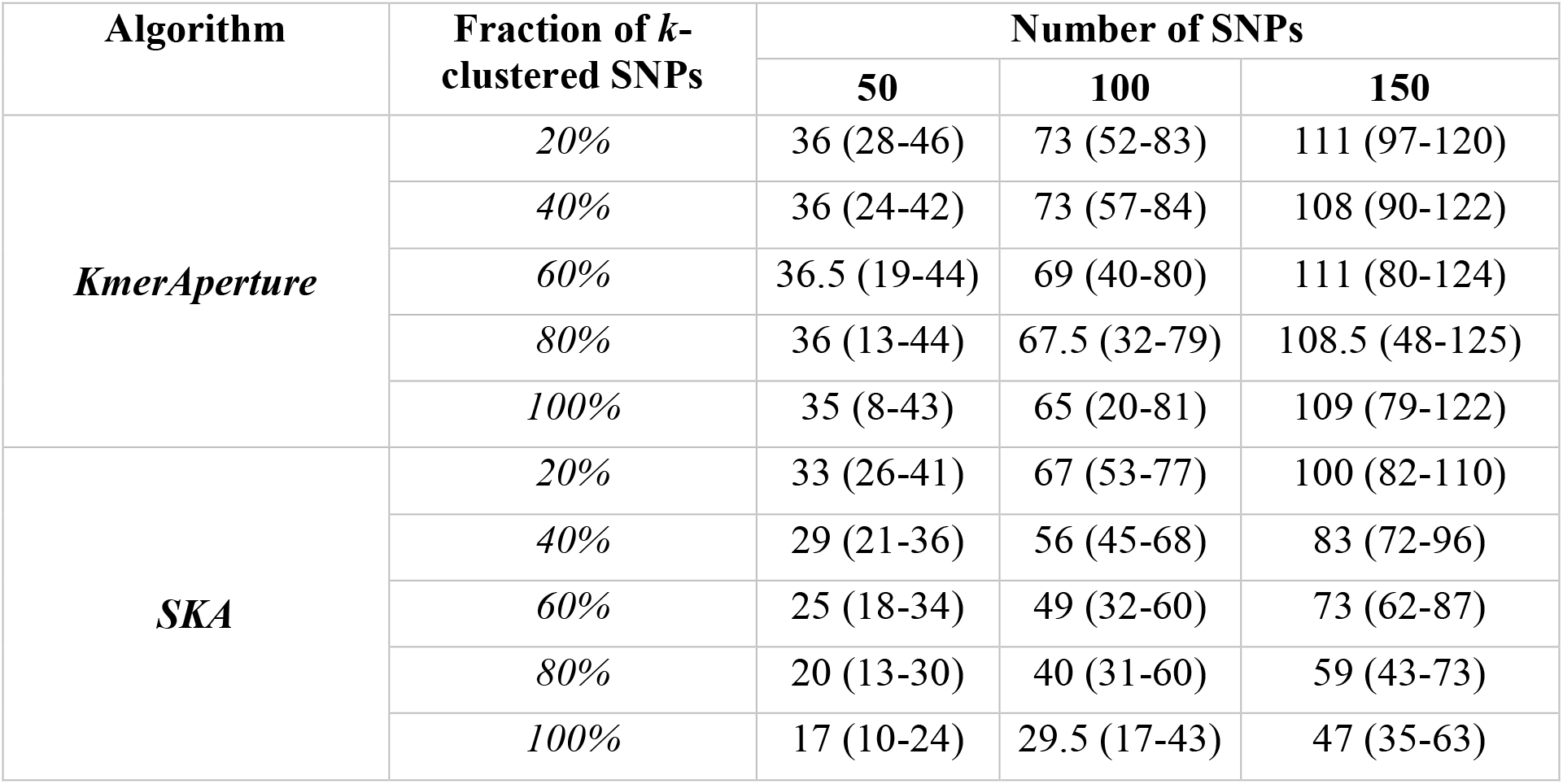
Performance by SNP retrieval for *KmerAperture* and SKA. There were three SNP amounts (50, 100, 150) and five fractions of *k*-clustering for each (20, 40, 60, 80 and 100% density). Within each category 50 genomes were simulated. The median retrieval is reported for each category with individual genome minimum and maximum retrieval

**Figure 2.**
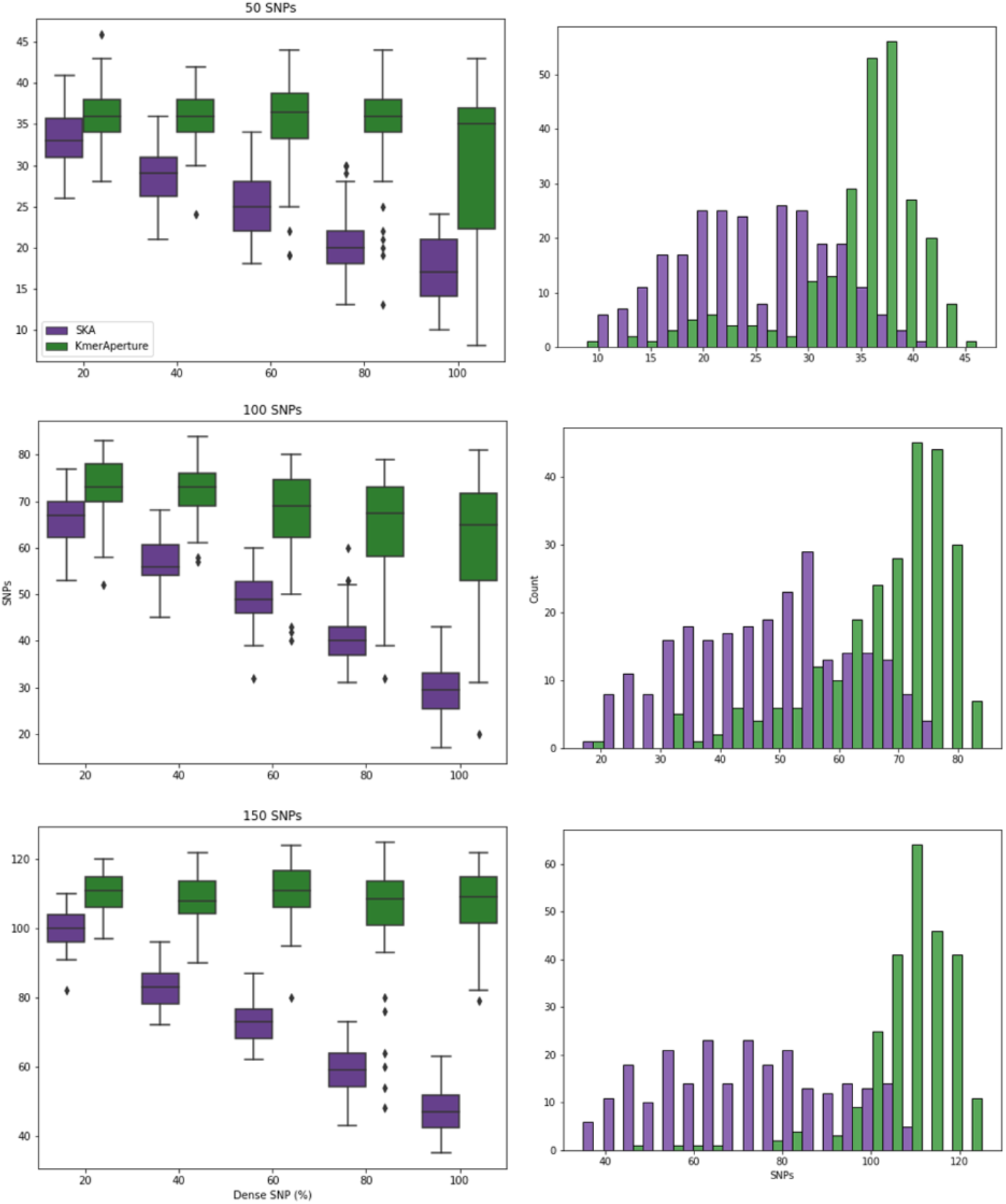
Boxplots (left) showing the distribution and median SNPs recovered by SKA and *KmerAperture*. Upper to lower are datasets with 50, 100 and 150 SNPs respectively. Each boxplot contains the values for 250 genomes, each with the respective number of SNPs but varying from 20% to 100% of SNPs that are within *k*=25 of one another. Histograms (right) show the distribution of SNPs recovered across all percent SNP *k*-clustering fractions

### Application to a cluster of *Escherichia coli* genomes

The genomes in the *E. coli* dataset were the most diverse analysed with a median (range) of 82 (29-870) SNPs relative to the reference and large accessory differences. Of 749 *E. coli* genomes 476 were <100 SNPs and 53 were <50 SNPs. By querying each *E. coli* genome against the reference, MCJCHV-1, *KmerAperture* predicted a median (range) of 84 (28-980), 85 (29-960) and 86 (29-892) SNPs for *k*=19, 25 and 31 respectively. SKA produced SNP estimates with a median (range) of 117 (54-465), 93 (38-398) and 84 (32-349) for *k*=19, 25 and 31 respectively. SKA trended towards overestimation in those with ≤150 SNPs and underestimation in those above (Figure 3; Figure S1). This is reflected in the SNP ranges and median estimations at *k*=19 and 25, with an improvement on median SNPs at *k*=31.

**Figure 3.**
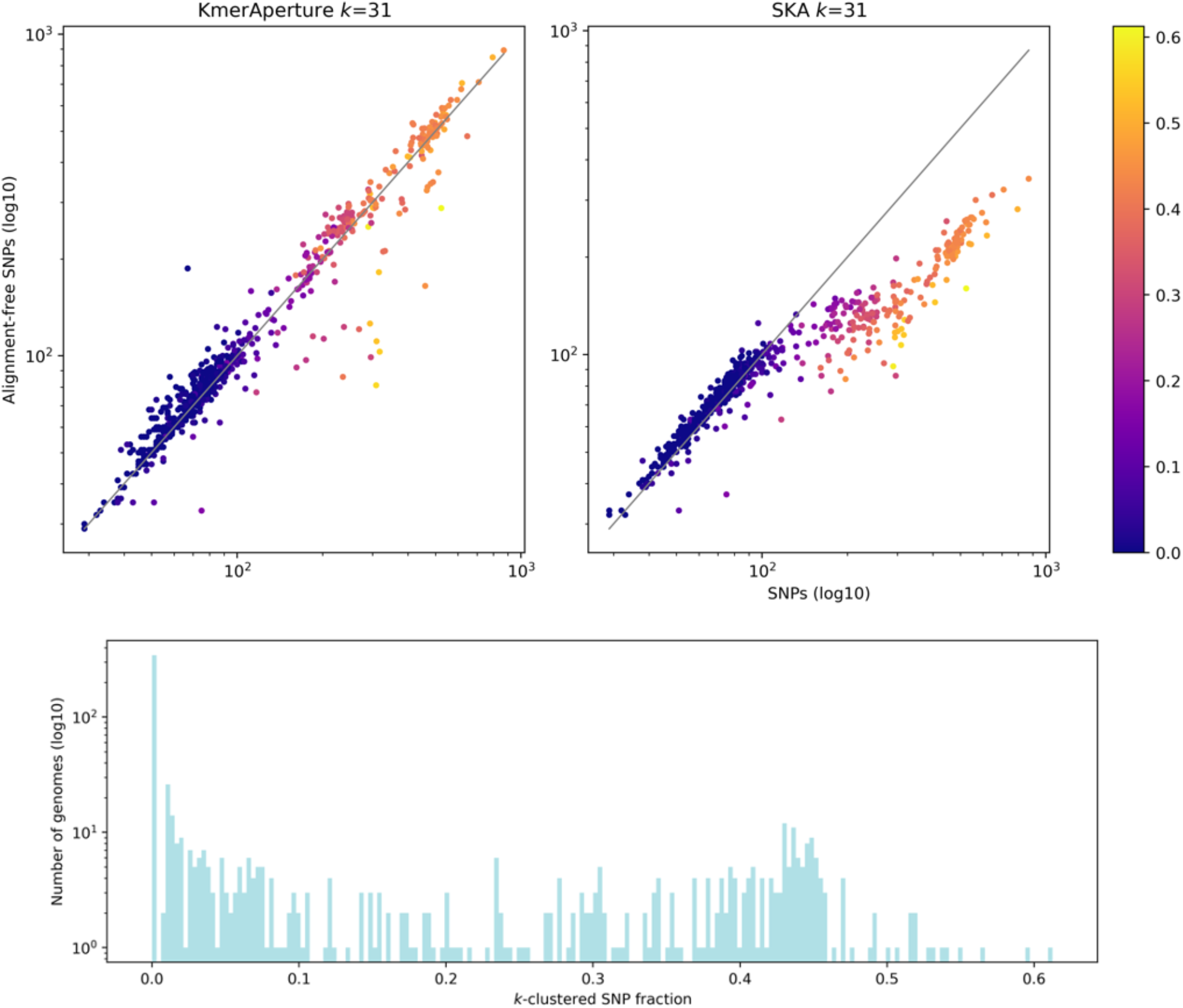
(Upper) scatterplots of *KmerAperture* (upper left) and SKA (upper right) SNPs at *k*=31 vs snippy SNPs. Data points are coloured by the fraction of SNPs that are *k*-clustered. Lower, a histogram of the number of genomes with that fraction of SNPs that are *k*-clustered with *k*=31

In this moderately diverse genome dataset, the SNPs extracted by *KmerAperture* fit the real SNP counts closely, according to the to the root-mean-squared error performance (RMSE) on log10 values. Crucially, the performance was largely independent of choice of *k*, with little difference across RMSE values. The RMSE with *KmerAperture* was 0.06 for *k*=19 and 0.07 for *k*=25 and 31. The level of under- and overprediction appeared to also be independent of real SNP count values. For comparison, SKA systematically overpredicted SNP counts when real values were ≤150 SNPs and underpredicted for >150 SNPs. The RMSE values were best for SKA at *k*=25 (0.15) followed by 0.17 for *k*=19 and 31. *KmerAperture* is designed to accurately extract SNPs within closely related genomes but the *E. coli* datasets shows good performance in those up to >400 SNPs (n=95 genomes). The RMSE values for those with >400 SNPs were 0.06 for *k*=19 and 31 and 0.08 for *k*=25 with *KmerAperture*, whereas SKA ranged from 0.28 to 0.34.

The spatial density of SNPs had a median of >*k* for each size of *k* with the *E. coli* genomes and the majority of genomes did have a single pair of SNPs <*k* apart. At *k*=19, 369/748 genomes had no SNPs within *k*, decreasing with *k*=25 and 31 to 356/748 and 344/748, respectively. However, significant proportions of SNPs in some genomes were *k*-clustered, i.e., within *k* of one another. For instance, at *k*=19, 19% (147/748) of the genomes had ≥25% of the SNPs being *k*-clustered, including one genome whose SNPs were 48% 19*-*clustered (i.e., with 99/207 couples of SNPs having distance ≤19). As *k* increases in size, the number of *k*-clustered chains also *k* increases. As such, the greatest proportion was recorded at *k*=31, whereby 27% (201/748) of genomes had ≥25% of SNPs being *k*-clustered. *KmerAperture* performance, however, was not reduced in the genomes having the highest SNP density (Figure 3).

The minimum reference genome bases not present in the respective query genomes based on MUMmer (unaligned sequences >60bp) was 273bp and maximum 920,525bp with a median of 47,186bp. *KmerAperture* recovered a per query genome median (range) of 46,952bp (98bp-914,782bp) with *k*=19 increasing with *k*=25 and 31 with a median (range) 48,746bp (210bp-921,213bp). The RMSE on log10 values reported a close fit for all values of *k* with 0.038 and 0.025 for *k*=19 and 25 respectively and the closest fit at *k*=31 with 0.023 (Figure S2).

### Application to a cluster of *Salmonella* Typhimurium genomes

The *Salmonella* Typhimurium dataset (n=1,264 genomes) represent a low diversity dataset with many highly similar genomes relative to the reference. The real SNP counts relative to the reference were median (range) 22 (1-164), with 1,176 genomes with <50 SNPs and 149 with ≤10 SNPs. *KmerAperture* was run for all genomes and extracted a median (range) of 21 (1-520), 21 (1-519) and 22 (1-526) SNPs for *k*=19, 25 and 31 respectively. Median (range) SNP counts with SKA at *k*=31 were similar to *KmerAperture* and the true figures, with 25 (2-136) SNPs. On the other hand, the results produced by SKA for *k*=19 and 25 were less accurate based on their summary statistics with median (range) 58 (19-195) and 35 (9-160) respectively

The SNP counts determined by *KmerAperture* fit the real SNP counts closely. There was similar performance with RMSE on log10 values of 0.07 for *k*=19 and 0.08 for *k*=25 and 31. For *k*=31 SKA and real SNP counts RMSE was similar to *KmerAperture* with 0.09. It was sensitive of choice of *k* however, producing an RMSE of 0.27 for *k*=25 and 0.48 for *k*=19. SKA produced a consistent level of SNP count overprediction along the range of SNP diversity. SKA performance was best at *k*=31, with overprediction minimised and a small amount of underprediction of the highest values (Figure 4).

**Figure 4.**
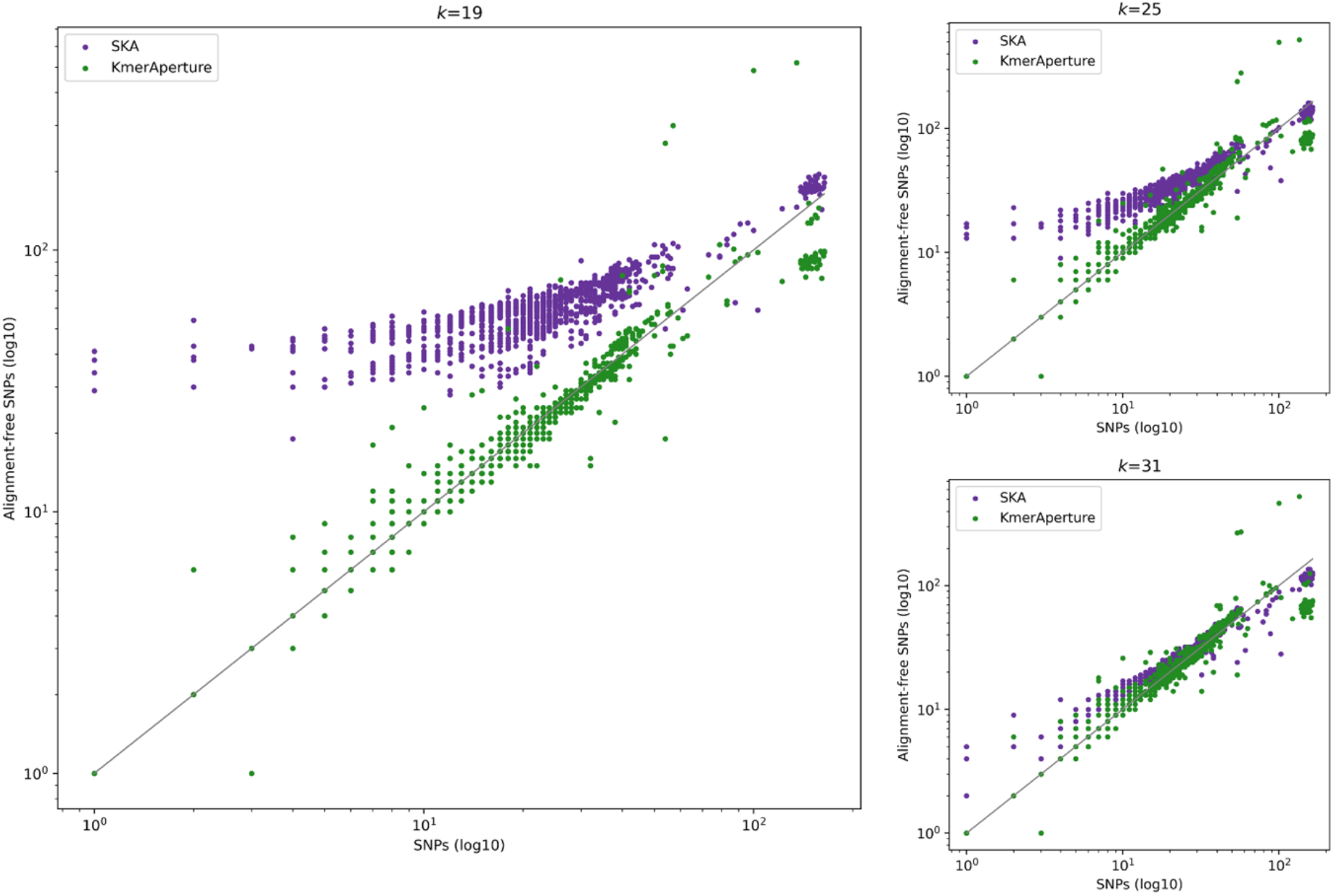
Results on *Salmonella* Typhimurium SNP counts. Scatterplot of *KmerAperture* and SKA SNP estimates vs real SNPs, for *k*=19, 25 and 31

A flexible core genome alignment was derived from the snippy, SKA and *KmerAperture* whole genome reference-anchored alignments and the pairwise SNP counts extracted. There were 798,216 exclusive pairs with a median (range) of 34 (0-822) pairwise cgSNPs. Graphs were constructed with edges between genomes (nodes) if they met a SNP threshold of ≤2, ≤5 or ≤10 SNPs. *KmerAperture* generated graphs with 646, 941 and 1187 nodes for ≤2, <5, ≤10 respectively. These were comprised of single linkage clusters (connected components). At ≤2 SNPs, *KmerAperture* generated 145 connected components, 72 of which contained at least 3 genomes, compared with 152 real SNP clusters at ≤2 SNPs including 76 with at least 3 genomes. There was similar recovery of clusters for ≤5 and ≤10 SNPs with 82 and 34 clusters with >2 genomes with *KmerAperture* respectively compared with 80 and 37 real SNP clusters. SKA reconstructed fewer SNP clusters (>2 genomes) with 38 at ≤2SNPs, 65 at ≤5 SNPs and a greater number than with real SNPs with 65 at ≤10 SNPs.

It was determined by MUMmer that query genomes did not align to a median (range) of 45,020bp (6,917bp-216,154bp) of the reference genome. The median estimate for *KmerAperture* ranged from 43,480bp to 46,925bp for *k*=19 and 31 respectively (Figure 5). The RMSE of the log10 values fit best with *k*=25 with 0.018 and fit closely for *k*=19 and 31 with 0.021 and 0.025 respectively. This represents recovery of the majority of invariant sites between query genomes and the reference. Additionally, as with SNP counts recovered by *KmerAperture* recovery of *aligned* reference sites is independent of *k* choice with the *S*. Typhimurium dataset.

**Figure 5.**
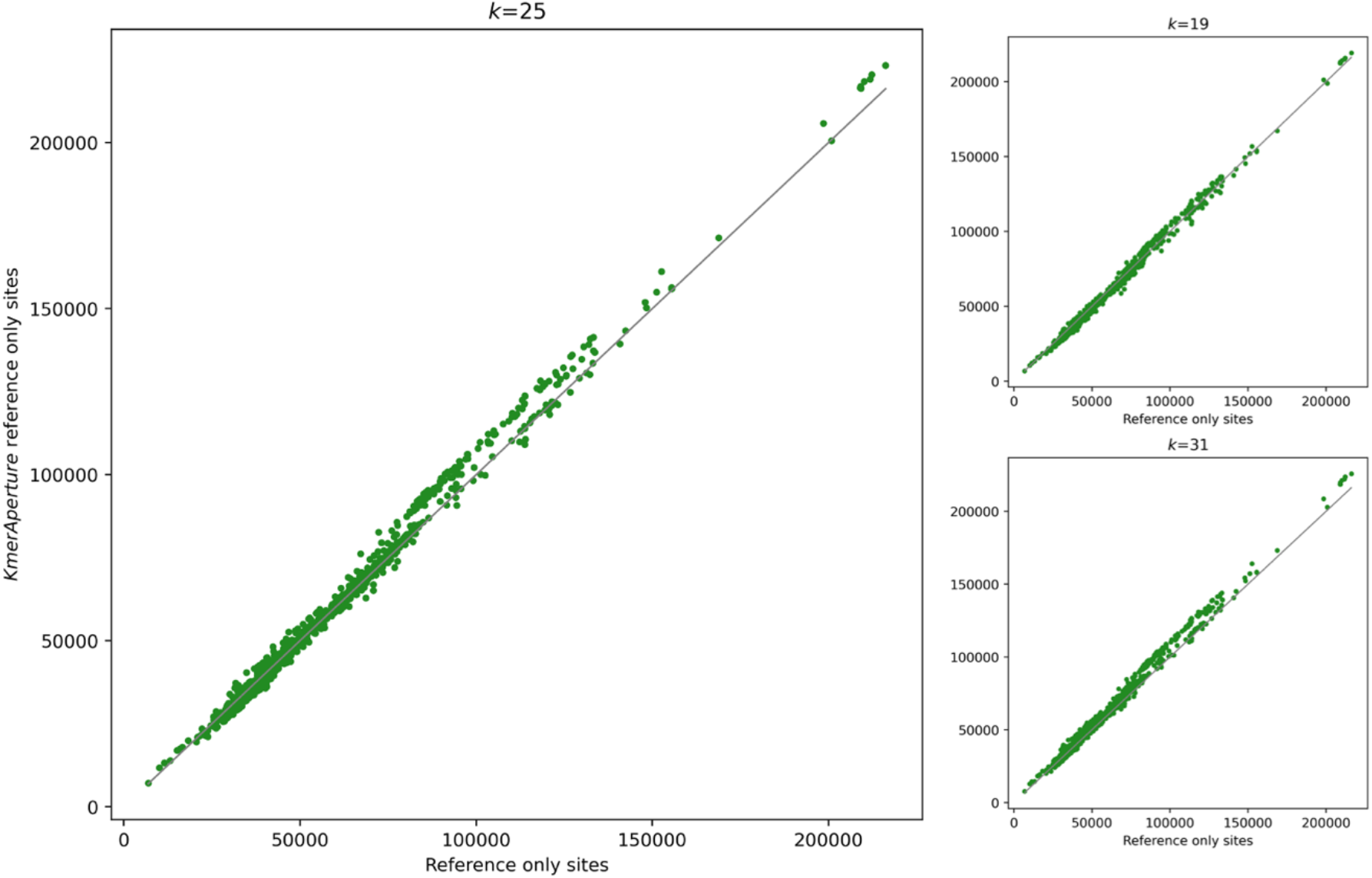
Results on *Salmonella* Typhimurium number of reference sites not present in query genomes determined by *KmerAperture* compared with unaligned reference bases with MUMmer, at *k*=19, 25 and 31

### *KmerAperture* runtime

The runtimes for *KmerAperture* and SKA were recorded as the mean per-genome runtime by dividing the overall runtime by the number of genomes. In all cases there was a slight increase in runtime with increase in size of *k*. Mean per genome runtime with the real genome dataseets was faster with *KmerAperture*. With *KmerAperture* runtime averaged between 10.41 and 13.87 seconds for the *Salmonella* Typhimurium genomes and between 13.57 and 14.82 for the *E. coli* genomes for *k*=19 to *k*=31 respectively. Whereas SKA averaged between 22.8 and 23.8 seconds for the *Salmonella* Typhimurium genomes and between 24.1 and 26.4 for the *E. coli* genomes for *k*=19 to *k*=31. Further with *KmerAperture*, query genomes may be analysed concurrently across multiple processors, making the time to extract an alignment from 1,256 *Salmonella* Typhimurium genomes (*k*=19) 21 minutes with 10 processors, or ∼1 second per genome.

## DISCUSSION

The *KmerAperture* algorithm is based on the few axioms we have of how relative sequence differences are represented by comparing *k*-mers. In sequences of the same length, with >*k* spaced SNP positions and no other variation, the number of unique *k*-mers / *k* will represent the number of variant positions. In reality, bacterial genomes have large unaligned regions, *k*-clustered SNPs, indels, structural rearrangements and other complexities. We demonstrated with simulated and real datasets that by retaining *k*-mer synteny we can accurately recover SNPs, including many of those that are *k*-clustered. Initially the algorithm filters mismatched *k*-mers by their original order, nominating contiguous series of length *k* as being generated by a single SNP difference between the genomes. This is followed, by matching the middle *k*-mers without the middle base. When several SNPs’ positions occur ≤*k* bp from each other, series of length >*k* are generated. By pattern matching between same length series, we explicitly consider this possibility. The persistent problem in *k*-mer based alignment-free analyses is here resolved. Essentially, *KmerAperture* foregoes alignment as matching flanking regions to *k*-mer series are implicit in their presence in the *k*-mer set intersection, which can be rapidly determined. With *KmerAperture*, the entire relative diversity of genomes is encoded in the syntenic *k*-mer series. Methods may be improved further to types of diversity such as indels and sequence rearrangements.

Since *KmerAperture* determines the position of each SNP, it can be used to generate a reference-anchored alignment containing all the SNPs found in a set of genomes of interest. Unlike a simple SNP matrix with no notion of SNP locations, this *KmerAperture* output can be to used to perform a recombination-aware phylogenetic inference, for example using ClonalFrameML^17^ or Gubbins^18^. In this context, it is especially important that *KmerAperture* attempts to reconstruct SNPs within *k* bp of each other, since bacterial homologous recombination, especially when the donor and recipient are distantly related, has a high potential to generate such clusters of SNPs^19^.

Unlike other tools that are focused on SNP detection, *KmerAperture* is also able to identify accessory regions, and may therefore be developed for pan-genome analyses^20^. It would also be possible to extract accessory sequence positions, prior to alignment. The computational and temporal challenge of alignment is related both to the number of sequence comparisons but also the diversity of the sequences under study. Reducing this diversity, especially in genome samples of unknown diversity, would reduce the cost of alignment. *KmerAperture* could also be adapted to progressively remove accessory regions as genomes are added, compare the genomes to a core genome alignment, or to a reference.

In the examples tested *KmerAperture* works as well in situations where genomes differ by <50 SNPs as those with >400. It could now be examined whether clustering on these SNPs, or construction of a phylogenetic tree from the alignment may be used as a scalable alternative to sub-MLST lineage discrimination. For instance, clustering the *S*. Typhimurium genomes by SNP thresholds suggest that this may be a viable tool to test on numerous species, to provide results comparable to alignment. This is crucial because whilst existing tools may be accurate once very similar genomes have been identified, this is the first alignment-free methodology able to discern clusters of very similar genomes.

The datasets were chosen to reflect real-life scenarios of the type of diversity according to which we might cluster genomes, either by genomic typing or by methods, such as MinHash, not involving alignment. Divergent genomes can be identified and removed, retaining only the low-diversity genomes to be analysed further, possibly using alignment-based comparison. In a scenario where there is a survey of bacterial genomes, for instance in a clinical setting, it is important to rapidly differentiate species, lineage, sub-lineage and possible outbreak clusters based on genetic distance. The size and diversity of these datasets can render alignment-based methods difficult and possibly intractable unless an excess amount of computational resources is used. *KmerAperture* runs in a few seconds on a laptop, and genomes may be compared concurrently, making it a scalable approach for genomic surveillance.

## METHODS

### Using synteny to distinguish core and accessory variation

The data for analysis consists of a single reference genome and several query genomes, all of which have been assembled *de novo* and may therefore be made of multiple contigs^21^. Both reference and query genomes are decomposed into *k*-mers and canonicalised, which involves reverse complementing each *k*-mer, comparing to the original and selecting the lesser *k*-mer. The *k*-mer window size (*k*) is defaulted to 19, whilst various other *k* sizes were also explored.

For each query genome the full set of *k*-mers is constructed for the query and reference (sets *R* and *Q* respectively). Next the relative complement of each set (*R*^*c*^ and *Q*^*c*^) are determined by subtracting the sets from one another. The original lists of *k*-mers (which unlike the sets are not deduplicated) were enumerated. The *k*-mers of the relative complements were then matched back to their respective *k*-mer lists to derive their synteny. Each matched *k*-mer position is then stored and sorted as ranges or *k*-mer ‘series’. The length of these series is then used to evaluate the underlying comparative diversity that has generated these mismatching *k*-mers.

Series of *k*-mers that equal *k* in length (*L*) are provisionally taken to be the result of a SNP, whilst those greater than *k* in length are stored as accessory/other. It was enforced that *k* must be an odd number so that the middle *k*-mer may be extracted from all series of length *k*, the middle base removed and then the number of *k*-mers that would match without the middle base counted.

Whether there are SNPs within *k* of one another is evaluated next, including accounting for the possibility of chaining of multiple SNPs within *k*. Series of length *L* ranging from *k*+2 to *S*(*k*-1)+1 have their original sequences reconstructed, where *S* is the maximum number of ‘chained’ SNPs (defaulted to 10). Unlike with single SNPs there isn’t an explicit relationship expected between *L* and number of polymorphisms. Instead, all possible pairs of the same *L* are compared including the respective reverse complement pairings, mismatching bases counted and a filter of ≤25% divergence applied. All mismatches must not be contiguous in the sequence, with spacing of at least 1bp required.

### Implementation and benchmarking of *KmerAperture*

*KmerAperture* is written in python 3 and depends on the python packages numpy^22^, biopython^23^ and the screed module of the khmer package^24^. An OCaml, compiled, parser processes input genomes in a computationally efficient manner, decomposing them into *k*-mers and canonicalising them. Visualisations were generated in python with the use of seaborn^25^ and matplotlib^26^ in the Jupyter Notebook environment^27^. Conda was used for package management^28^. *KmerAperture* was compared with SKA *map* SNP estimations. *KmerAperture* was run on assembled genomes across a range of values of *k* (*k*=19, 25 and 31), where *k* must be an odd number. SKA requires that the *k* value be generated by an integer divisible by 3, that represents the length of each sequence flanking a flexible base. As such, a range of flanking lengths was selected of 9, 12, and 15 to generate the same *k*=19, 25, and 31 as in *KmerAperture*. Real genome SNP counts were derived from the tool snippy^29^, whilst unshared sequences were determined by MUMmer dnadiff^30^.

### Data acquisition and processing

The reference genome *Klebsiella pneumoniae* HS11286 (NC_016845)^31^ was downloaded and used as the template for each simulated genome. Two sets of simulated genomes were constructed: in the first set genomes relative to the reference have additional sequence and SNPs and in the second set all genomes are the same size as the reference with SNPs only.

For the first set, 500 genomes were simulated with between 1-100 SNPs and 1-30 accessory sequences of 1,000-10,000bp. The number of accessory sequences, the location where they were inserted and the genome positions to mutate, were all uniformly sampled at random whilst the number of SNPs were sampled from a log10 distribution. There was no relationship simulated between amount of accessory sequence and number of SNPs. All genomes were randomly assigned between 1,000 and 300,000bp of accessory sequence.

For the second set, genomes were simulated with SNPs relative to the reference only. Five subsets, each with 100 genomes were constructed. All genomes had 50 SNPs relative to the reference. The five subsets were constructed to contain 20%, 40%, 60%, 80% and 100% of the SNPs (n=10,20,30,40,50 SNPs respectively) to be within *k* of at least one other SNP. For each genome it was randomly determined how many SNPs, up to 10, would be within *k* and how large the spacing between each SNP was, up to *k*. The size of *k* was fixed to 25.

Real genome datasets were also used to assess *KmerAperture* performance, a low diversity and medium diversity within-lineage population. The first was a medium diversity cluster of genomes, subset of the pandemic *Escherichia coli*, MLST-defined cluster ST1193. The ST1193 population records were further subset to the HierCC^32^ HC20-level cluster 571 (n=1,331), where they were filtered for those with NCBI ‘biosample’ identifiers (n=1,233), whereby 757 assemblies were successfully downloaded. Finally, 8 genomes were removed for being too diverse (>1,000 SNPs including 6 at >30,000) resulting in 749 for analysis. Within cluster HC20 level cluster 571, reference genome *E. coli* MCJCHV-1 sequence (chromosomal: CP030111.1), was selected for mapping and variant calling. EnteroBase^33^ was accessed and searched for these criteria on 22/02/2023.

The second cluster of real genomes was a subset of the *Salmonella enterica* subsp. *enterica* Serovar Typhimurium (*Salmonella* Typhimurium) MLST defined cluster 34 (ST34). The ST34 population records were further subset to the HierCC^32^ HC5-level cluster 302 (HC302, n=2,399). Within cluster HC302 cluster, the completely assembled genome *S*. Typhimurium S04698-09, was selected as the reference. EnteroBase^33^ was accessed and searched for these criteria on 22/02/2023. All genomes with NCBI biosamples were selected for download (n=1,768), of which 1,283 successfully downloaded. A further, 3 genomes were excluded due to being too diverse (>300 SNPs) whilst 16 were removed for containing non-{ATCGN} bases. Finally, 8 genomes were removed for reporting an MLST other than ST34, resulting in the final set of 1,256 genomes.

All genomes were downloaded from the NCBI genomes database (Supplementary Table 1).

## DATA AVAILABILITY

All genome data is publicly available and acquired from the National Centre for Biotechnology Information (NCBI) databases. All accession IDs are provided in supplementary information. Open-source code is available from: https://github.com/moorembioinfo/KmerAperture

## AUTHOR CONTRIBUTIONS

All authors contributed to this research and resulting manuscript. Their contributions are listed here in accordance with the Contributor Roles Taxonomy (CRediT). ***Matthew P. Moore:*** conceptualisation, methodology, software, validation, formal analysis, data curation, writing - original draft, visualisation. ***Mirjam Laager:*** conceptualisation, methodology. ***Paolo Ribeca:*** methodology, writing - review and editing, supervision. ***Xavier Didelot:*** conceptualisation, methodology, writing - review and editing, resources, supervision, project administration, funding acquisition.

## ACKNOWLEDGEMENTS

This study is funded by the National Institute for Health and Care Research (NIHR) Health Protection Research Unit in Gastrointestinal Infections, a partnership between the UK Health Security Agency, the University of Liverpool and the University of Warwick. The views expressed are those of the author(s) and not necessarily those of the NIHR, the UK Health Security Agency or the Department of Health and Social Care.

## SUPPLEMENTARY FIGURE LEGENDS

**Figure S1.** Results on *E. coli* ST1193 SNP distances. Scatterplot of *KmerAperture* and SKA SNP estimates vs real SNPs, for *k*=19, 25 and 31.

**Figure S2.** Results on *E. coli* ST1193 number of reference sites not present in query genomes determined by KmerAperture compared with unaligned reference bases with MUMmer, at *k*=19, 25 and 31.

