## Supplementary figures and images for "*KmerAperture*: Retaining *k*-mer synteny for alignment-free extraction of core and accessory differences between bacterial genomes"

### SI1

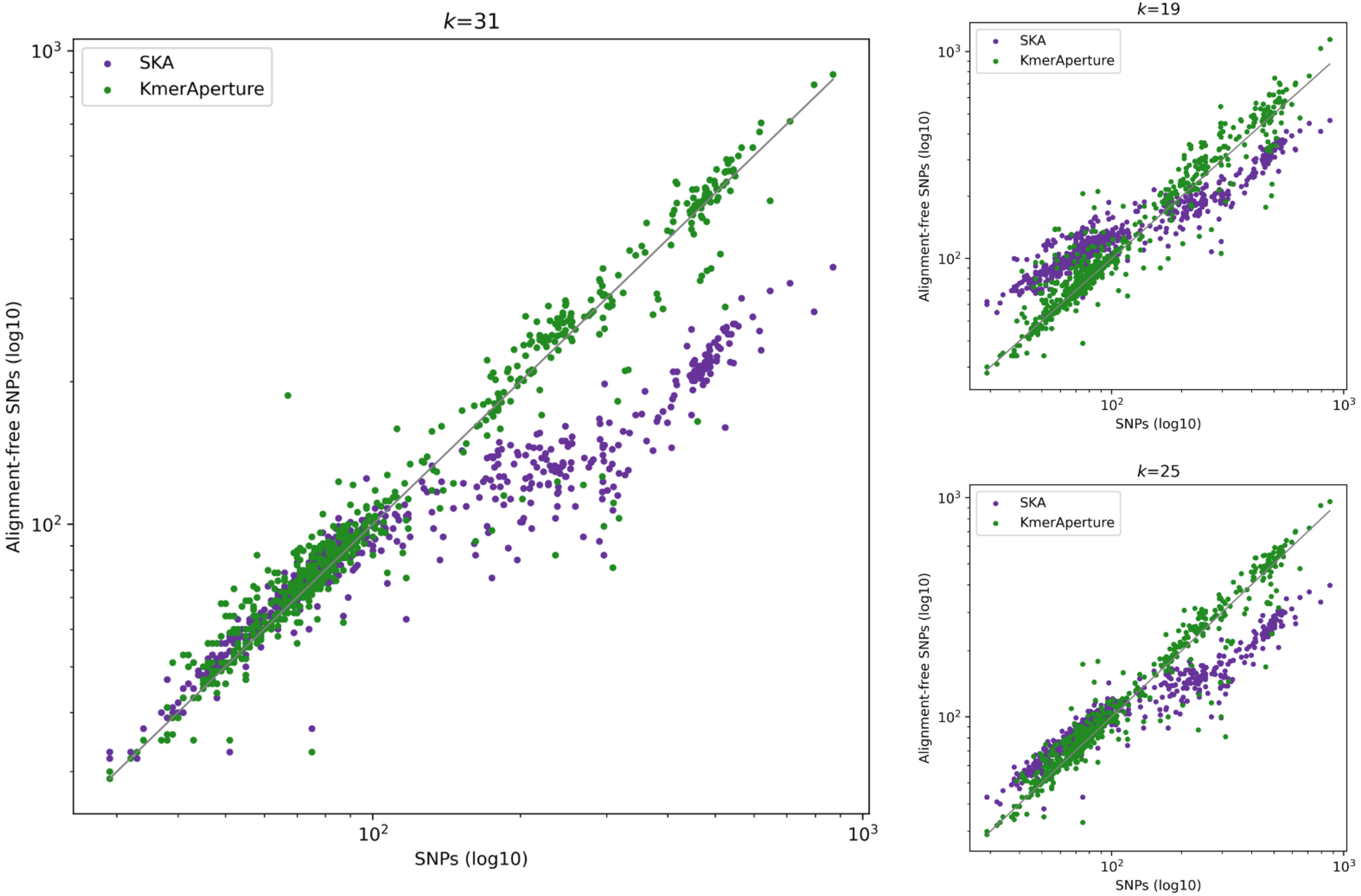

### SI2

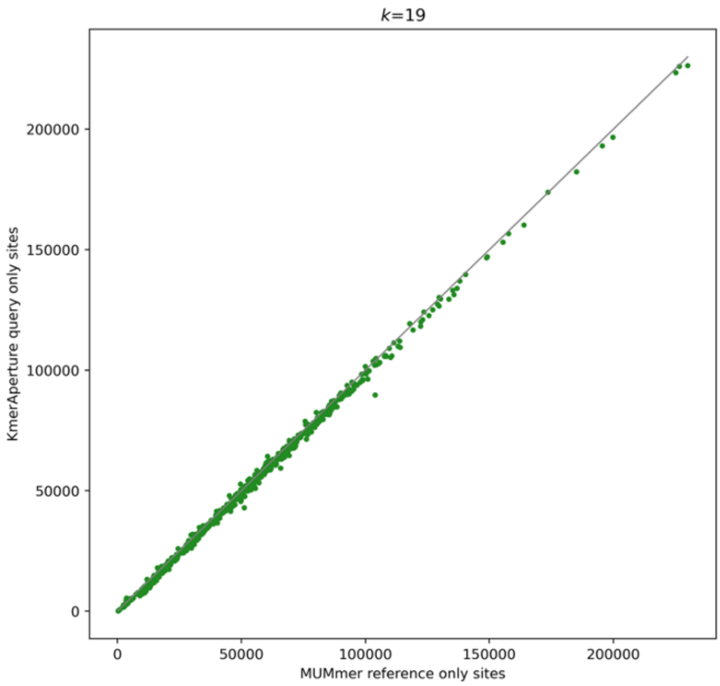

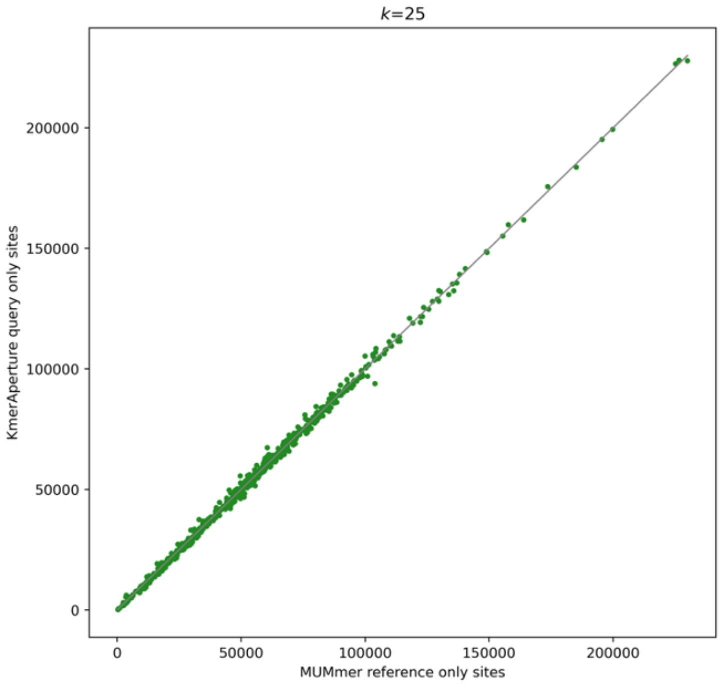

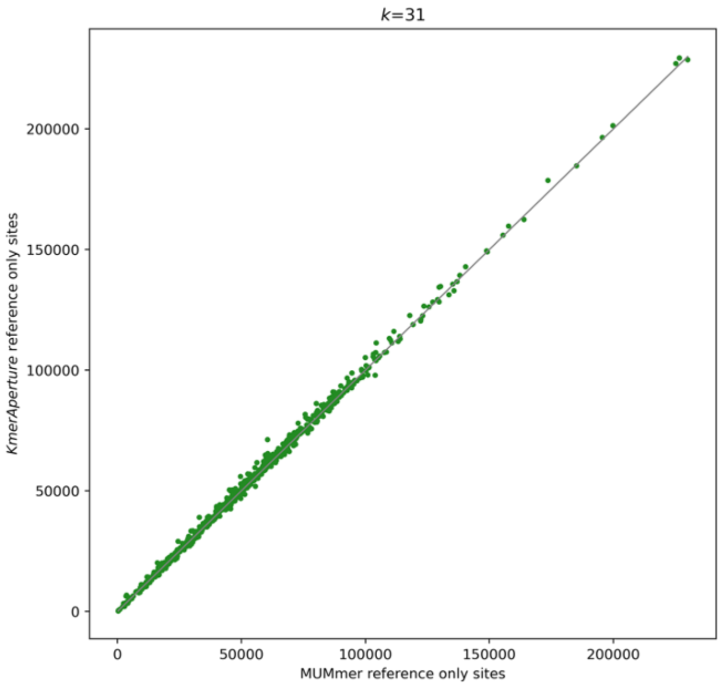
